# QuNex Recipes: Executable, Human-Readable Workflows for Reproducible Neuroimaging Research

**DOI:** 10.1101/2025.11.08.687330

**Authors:** Jure Demšar, Aleksij Kraljič, Andraž Matkovič, Samuel Brege, Lining Pan, Zailyn Tamayo, Clara Fonteneau, Markus Helmer, Jie Lisa Ji, Alan Anticevic, Cole Korponay, Melissa Salavrakos, Daniel Drucker, Matthew F. Glasser, Lisa D. Nickerson, Youngsun T. Cho, Grega Repovš

## Abstract

Preprocessing and analysis of neuroimaging data are technically demanding, often requiring a combination of multiple software tools, modality-specific pipelines, and extensive parameter tuning to match dataset characteristics. These complexities make it difficult to document workflows in sufficient detail to ensure complete transparency and reproducibility. To address these challenges, we introduce QuNex recipes, a framework for defining and executing complete neuroimaging workflows – encompassing data onboarding, preprocessing, and analysis – in a transparent, machine- and human-readable format. Recipes are implemented as an integrated feature of the Quantitative Neuroimaging Environment & Toolbox (QuNex), a containerized, open-source platform for end-to-end multimodal and multi-species neuroimaging processing. The recipes framework enables seamless integration of QuNex commands with custom scripts and external tools, capturing every processing step and parameter setting. A fully reproducible study can thus be shared and replicated by providing only (a) the QuNex version used, (b) the recipe file, and (c) the data. This approach standardizes workflow specification, enhances transparency, and enables one-command replication of complex neuroimaging analyses. By providing a standardized way to describe and share workflows, recipes facilitate open exchange of best practices and reproducible methods within the neuroimaging community.

## 1. Introduction

*QuNex (The Quantitative Neuroimaging Environment & Toolbox)* is an open-source software suite that provides an extensible framework for data organization, quality control, preprocessing, and analysis across multiple neuroimaging modalities and species (1). Since its release in 2023, more than 400 institutions and individual users have registered for access. To date, QuNex has been successfully used to process tens of thousands of scans across clinical and large-scale research datasets (2–7), including serving as the primary processing tool for the Human Connectome Project (HCP) team at Washington University for processing most of their recent data releases. QuNex is the only integrated neuroimaging platform that natively incorporates the HCP Minimal Preprocessing Pipelines (8) within a containerised framework that can be easily deployed in a high-performance computing (HPC) environment, enabling reproducible, scalable, and multimodal analyses across human and non-human species.

The original motivation for developing QuNex stemmed from persistent challenges in neuroimaging related to **reproducibility, transparency, and workflow integration** (9–12). In recent years, the neuroimaging community has advanced major initiatives to improve reproducibility, including the **FAIR data principles** (13), **the ReproNim** framework for reproducible neuroimaging (14), and efforts to standardize data and result representations through the **Brain Imaging Data Structure (BIDS)** (15), the **Neuroimaging Data Model (NIDM)** (16), and **DataLad** for distributed data and code management (17). BIDS Apps (18) improve reproducibility further by standardising the interface of individual containerised tools and defining a common input/output convention for BIDS-formatted datasets. These initiatives have substantially improved how neuroimaging data are organized and shared, yet the reproducibility of *analytic workflows* – the series of steps and parameter choices that transform raw data into scientific results – remains a major bottleneck.

Processing pipelines typically integrate multiple tools (e.g., Connectome Workbench, FreeSurfer, FSL, AFNI, SPM…), each with distinct dependencies and parameter sets. Even when executed within containers that guarantee software version consistency (19, 20), the absence of a unified, human-readable record of the workflow configuration often prevents exact replication. Although several tools address aspects of reproducible neuroimaging research, each has distinct limitations (Table 1). Preprocessing-focused tools like NiPreps (fMRIPrep, sMRIPrep, dMRIPrep) and qsiprep, which are built on the Nipype workflow framework (21), provide robust, containerised pipelines but are limited to preprocessing stages and are not designed for integrating custom postprocessing analyses within the same workflow (22, 23). Domain-specific platforms such as C-PAC and HALFpipe support end-to-end resting-state workflows but lack flexibility for external tool integration (24, 25). Nipoppy (26) takes a complementary approach, providing a lightweight protocol and Python framework that standardises dataset organisation and tracks participants through successive processing stages, though individual stages must be invoked separately and full analytic workflows are not expressed in a single specification. A related orchestration tool, BABS (27) automates the reproducible application of individual BIDS Apps across large datasets on HPC clusters using DataLad-based provenance tracking, but operates on one BIDS App per project and cannot compose multi-stage workflows in a single specification. MATLAB-based solutions (DPABI/DPARSF, CONN) provide end-to-end workflows with graphical interfaces, but use binary .mat files that limit version control integration, transparency, and human readability (28, 29). Brainlife (30) is a cloud-based platform that hosts a registry of community-contributed Apps with automated provenance capture. However, Brainlife’s App ecosystem is deliberately atomic (most Apps perform a single processing step), so end-to-end workflows must be assembled by users rather than invoked by a portable, version-controllable, specification file.

**Table 1.**
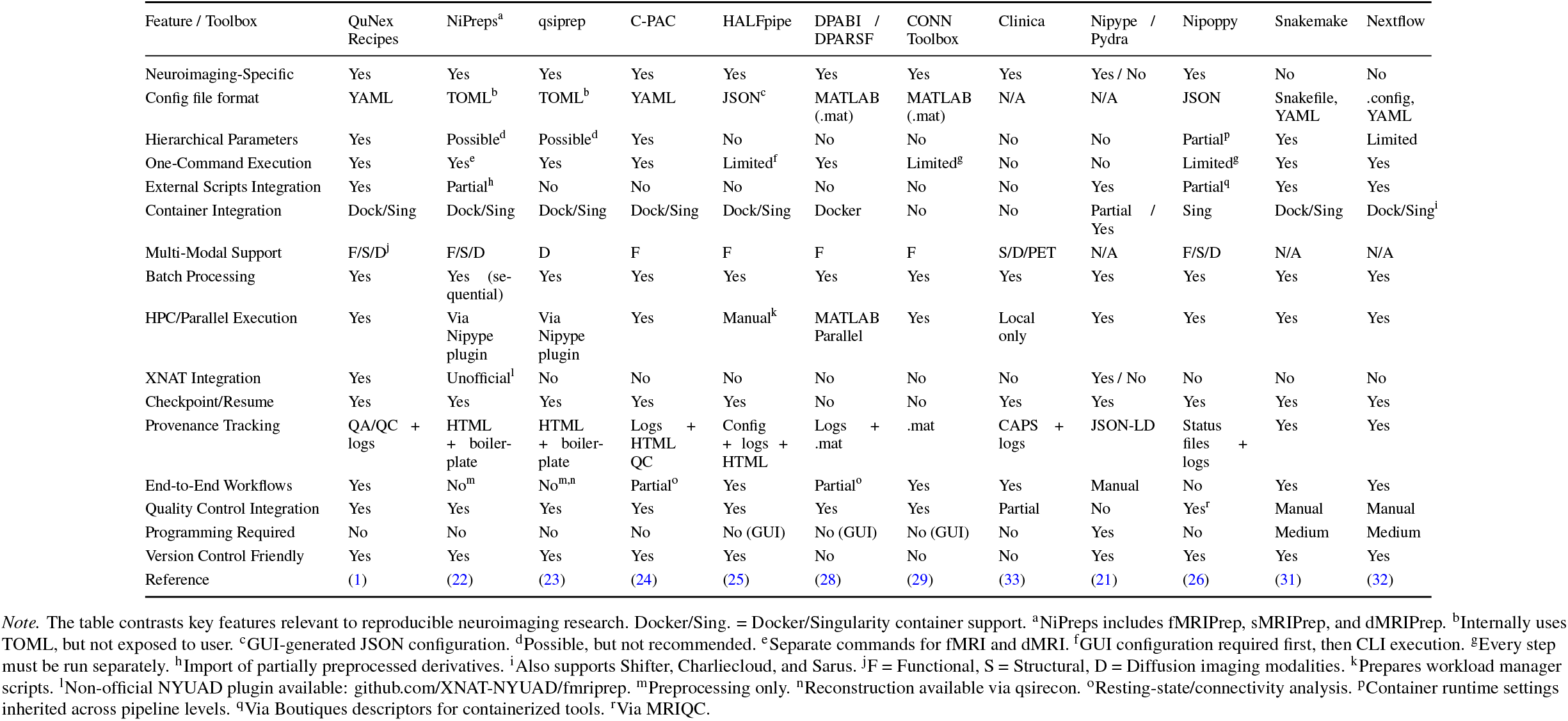
Comparison of QuNex Recipes with Existing Workflow Management Tools.

More general workflow managers such as Nipype (21), its successor Pydra, Snakemake (31) and Nextflow (32) provide flexible automation and provenance tracking. However, they require specialised scripting expertise and lack predefined, end-to-end neuroimaging pipelines and standardised quality control. Additionally, Snakemake and Nextflow lack native support for neuroimaging data organisation standards (BIDS). As a result, reproducing or adapting another group’s analysis often involves extensive manual reconstruction of undocumented steps and parameters, limiting openness and collaboration.

To directly address these challenges, we developed **QuNex recipes**, a framework that enables fully documented, reproducible definition and execution of neuroimaging workflows within the QuNex ecosystem. QuNex recipes are version control-friendly and fully transparent human-readable YAML files that define whole neuroimaging processing and analytics workflows and their configuration. These recipes are then automatically parsed by QuNex and unwrapped into a chain of commands that do the desired work. Unlike preprocessing-focused tools, QuNex recipes support end-to-end workflows from data onboarding through preprocessing and analysis. In contrast to most neuroimaging platforms, it enables seamless external script integration at any workflow stage, not just as pre- or post-processing hooks. It provides native XNAT integration for data management and a four-level hierarchical parameter system for flexible configuration. Importantly, QuNex recipes achieve this functionality without requiring programming expertise. The recipe specification file defines all processing and analysis stages (from data onboarding, preprocessing, and analysis, to quality control) along with the sequence of operations, and exact parameter values. The use of a recipe file within the new QuNex `run_recipe` command allows every step of a preprocessing and analytic workflow to be encoded in a single, shareable file. Note that the main functionality of QuNex recipes is to seamlessly link QuNex processing, quality control and analytic commands together in a fully transparent and reproducible way. For a full list of supported commands and parameters, please see the official manuscript (1) or the the official documentation (https://qunex.readthedocs.io). Although recipes allow users to incorporate external commands and scripts within QuNex commands, this functionality is primarily intended for short utility scripts (e.g. downloading, preparing and transforming data) rather than for implementing or integrating custom, comprehensive neuroimaging pipelines. While this is possible, there are other tools that are better suited to this task (e.g. 31, 32).

As mentioned previously, the recipe framework is primarily designed to simplify the processing workflow by chaining together QuNex commands. While the external scripts framework is intended for executing simpler tasks and sharing relevant processing information for scientific publications and project replicability, it is possible to expand QuNex and integrate extensive custom workflows and pipelines into the recipes framework. This can be achieved either by using the QuNex extensions framework or by upgrading the container and installing extra tools and packages. The QuNex Docker container, for example, is public and can be used as a starting point in your Dockerfile, which you can then upgrade to accommodate your processing or analysis needs.

When used together with QuNex’s existing batch processing capabilities, QuNex recipes achieve three key goals: (1) **automatic documentation** of all processing and analytic steps within a single file; (2) **flexible aggregation** of diverse preprocessing and analytic workflows within a unified framework; and (3) **shareability and reusability**, allowing collaborators or external researchers to reproduce the full pipeline by providing only (a) the QuNex version used, (b) the recipe file, and (c) the input data.

This design bridges the gap between workflow automation and reproducible reporting, enabling transparent, one-command replication of complex multimodal analyses. By extending the principles of the FAIR and ReproNim movements to executable workflows, QuNex recipes provide a practical, scalable mechanism for open, collaborative, and fully reproducible neuroimaging research.

## 2. Methods

This section describes the design, and structure of the QuNex recipe framework. We first outline the format used to define recipes and explain how recipes specify each step of a neuroimaging workflow in a transparent and reproducible manner. Next, we detail the process of executing recipes within the containerized QuNex environment, followed by an overview of the internal workflow management system and its integration with high-performance computing resources. Finally, we describe how recipe-based processing can be facilitated for execution within the XNAT (Extensible Neuroimaging Archive Toolkit) environment.

### 2.1 QuNex recipes

QuNex processing recipes are defined using the YAML format (YAML Ain’t Markup Language; https://yaml.org). YAML is a compact and flexible human-readable data serialization standard that can be easily parsed by most programming languages. We selected YAML because it provides an intuitive and transparent structure for defining workflow specifications, while remaining compact and machine-readable. This design enables both non-programmers and advanced users to construct and interpret recipes without additional syntactic overhead.

Figure 1 illustrates the conceptual flow of a QuNex recipe workflow. A recipe defines all stages of neuroimaging processing – from data onboarding through structural and functional preprocessing to post-processing and analysis – within a single YAML file. Executed inside a versioned QuNex container, the recipe runs all specified commands, quality control steps, and optional external scripts or programs that can be integrated at any stage. Recipes can also be executed in parallel across sessions, enabling scalable processing on local or high-performance compute systems. Together with the data and QuNex version, the recipe forms a complete reproducibility bundle that enables one-command replication of the entire processing and analysis pipeline.

**Fig. 1.**
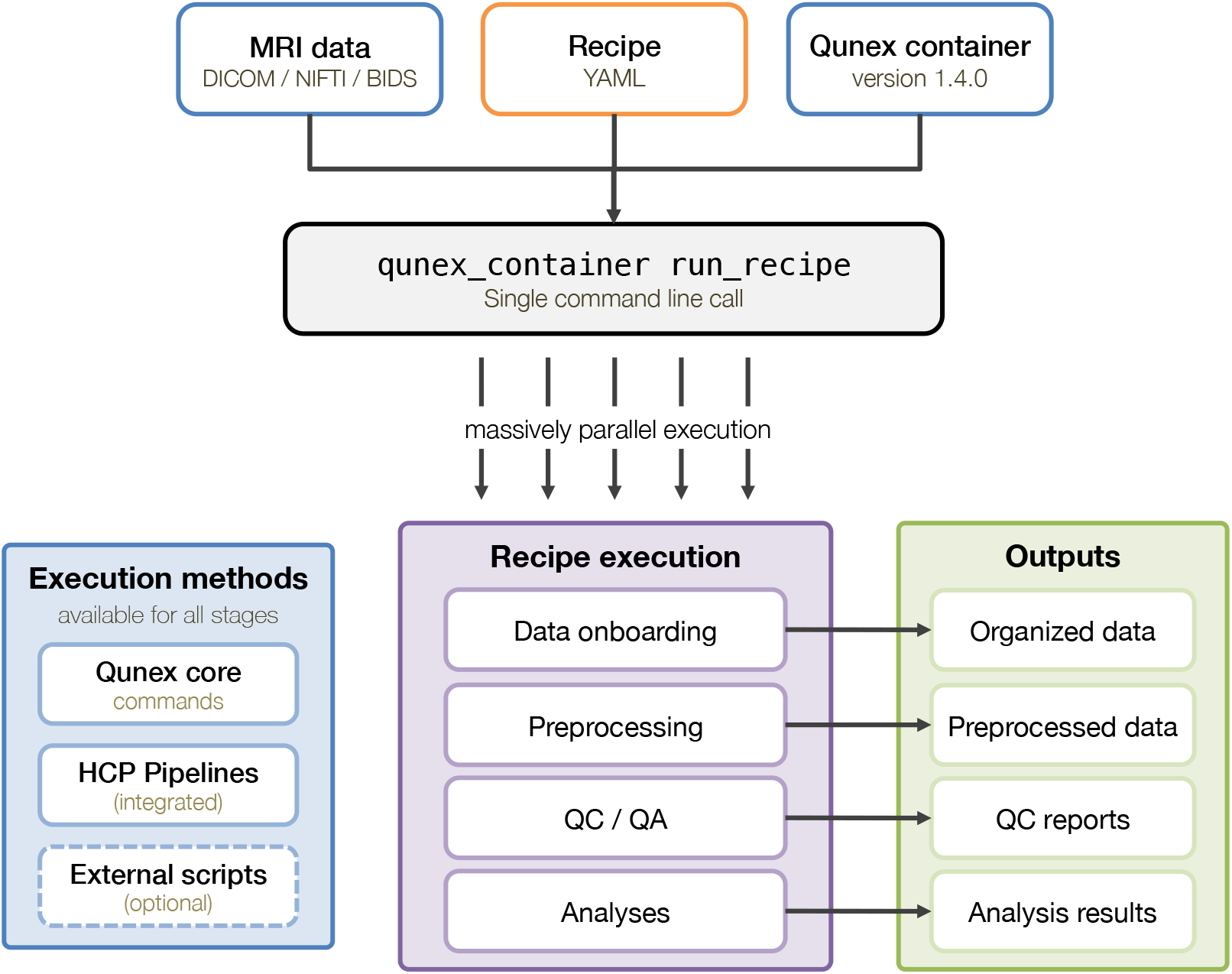
Schematic overview of a QuNex recipe structure. Each recipe file consists of optional global_parameters and one or more defined recipes. Within each recipe, commands are executed sequentially, with parameters inherited hierarchically from global, recipe, or command levels.

A QuNex recipe contains two main blocks at the root level of the YAML file: `global_parameters` and `recipes`. The `global_parameters` block is optional and defines parameters that apply across all commands listed within the `recipes` section. The `recipes` block specifies one or more processing or analysis workflows, each defined as a sequence of commands to be executed.

Each entry within the `recipes` block represents an individual recipe identified by name. For example, the code snippet in Listing 1 contains two recipes, `first_recipe` and `second_recipe`. A recipe may begin with an optional definition of *recipe-level parameters*, which apply only to commands within that recipe. The mandatory `commands` block defines the ordered sequence of commands to be executed and specifies command-specific parameters. For full flexibility purposes, parameters can be defined at four distinct hierarchical levels:

- command line call parameters (highest priority),
- command parameters,
- recipe parameters,
- global parameters (lowest priority).

Higher-level parameters take precedence over lower-level ones. When a parameter is defined at multiple levels, the value with the highest precedence is used in the final command call. Parameter values can be provided either statically or dynamically. Dynamic values are injected at runtime through the use of *mustache* notation (double curly braces, `{{…}}`), which allows substitution from operating system environment variables or external sources. This mechanism supports flexible integration of user-specific or system-dependent values while preserving full reproducibility of workflow specifications.

Listing 1 illustrates the logic and syntax of a typical QuNex recipe, showing the hierarchical organization of parameters, sequential command execution, and invocation of both internal and external processing routines.

**Listing 1.**
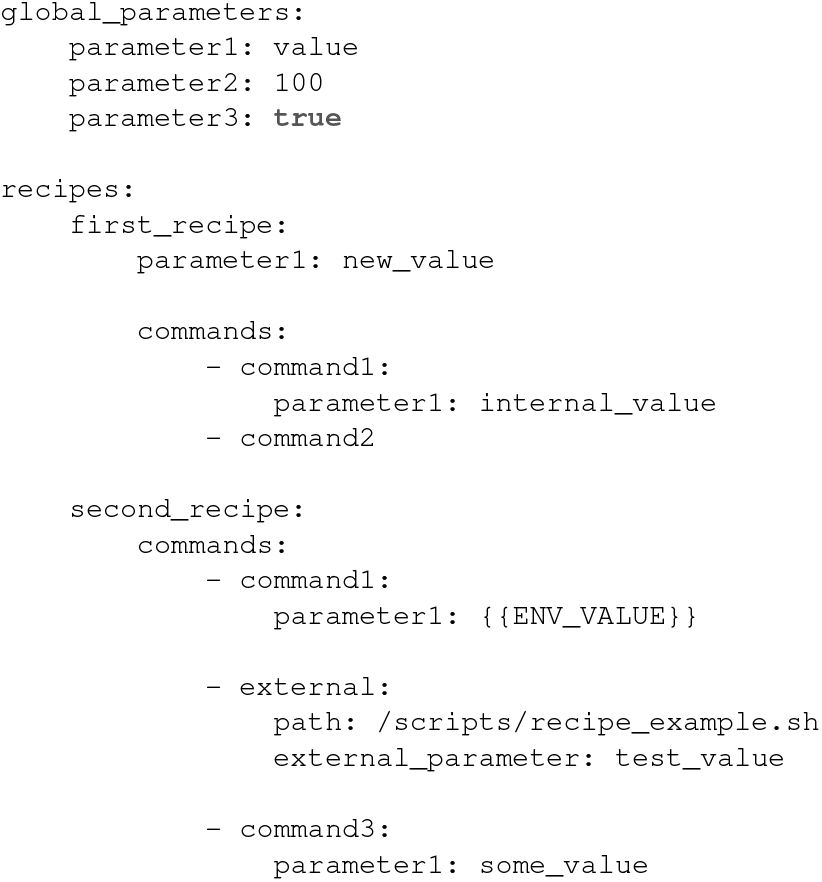
Example of a QuNex recipe file.

When executing `command1` from `first_recipe` the value `internal_value` will be used for `parameter1`. This is because the value defined at the command level (`internal_value`) has a higher priority than the value defined at the recipe level (`new_value`) and the global level (`value`). If `parameter1` value is provided with the command line interface call (see the next section) then that value will be used,since command line call parameters have the highest priority.

### 2.2 Executing a QuNex recipe

To execute a QuNex recipe (e.g., first_recipe) that is locally stored at `/data/qx_recipes/recipe.yaml`, the user can invoke the `run_recipe` command by running the following in a terminal:

~~~
qunex_container run_recipe \
  --recipe_file=“/data/qx_recipes/recipe.yaml” \
  --recipe=“first_recipe” \
  --container=“/data/qx_containers/qunex_suite-1.4.3.sif”
~~~

This command executes the sequence of QuNex commands defined in the first_recipe using the Singularity/Apptainer container with QuNex version 1.4.3 (the latest version at the time of writing). Additional details about QuNex containerization are available in the official documentation (https://qunex.readthedocs.io).

In this example, command1 is executed with `parameter1` equal to the `internal_value`, as this is the highest-precedence value defined for this parameter. If a parameter is additionally specified on the command line (e.g., `-parameter1=“value_x”`), that value takes priority, since command-line arguments override all lower-level definitions. Parameters `parameter2` and `parameter3`, defined under `global_parameters`, are automatically propagated to all commands. Commands that do not utilize a given parameter simply ignore it.

When executing the `second_recipe`, an environment variable must be defined for `ENV_VALUE`, which is dynamically injected into `parameter1` of `command3` through the mustache notation (`{{…}}`). Environment variables can be set directly through the `-bash_post` argument, which executes a short shell preamble after (post) entering the container, but before the recipe is launched:

~~~
qunex_container run_recipe \
  --recipe_file=“/data/qx_recipes/recipe.yaml” \
  --recipe=“second_recipe” \
  --bash_post=“export ENV_VALUE=value_injection” \
  --container=“/data/qx_containers/qunex_suite-1.4.3.sif”
~~~

The `second_recipe` also illustrates how external scripts, programs, or pipelines can be incorporated into a processing sequence. In this example, QuNex executes the following command as the second processing step:

~~~
/scripts/recipe_example.sh --external_parameter=“test_value”
~~~

By default, parameters of external scripts are prefixed with `-` and the separator between parameter name/key and value is the equal sign (=). However, since some scripts use a different syntax here, you can customise this through the `external_parameter_prefix` and `external_parameter_delimiter` parameters of the external block in the recipe. For example, setting the prefix to “-” and delimiter to “ “ will give the above parameter in the format `-external_parameter “test_value”`.

In addition to Bash (`.sh`) scripts, the recipe framework supports execution of Python and R scripts, as well as precompiled binaries. This flexibility allows integration of custom user code or third-party tools directly within QuNex workflows, while preserving full provenance and reproducibility.

The `qunex_container` script serves as a unified entry point for executing QuNex commands and recipes. It abstracts interactions with containerization and high-performance computing environments, simplifying workflow execution for users with varying levels of technical expertise and experience with containers and scheduling. Comprehensive documentation of its parameters and capabilities is provided in the original QuNex manuscript (1) and in the online documentation (https://qunex.readthedocs.io).

When executing commands through the recipe framework, QuNex will generate logs with three different levels of granularity. The highest level log is the recipe log, which includes the top-level information about the processing of the recipe as a whole. For each command in the recipe, the exact command call is printed after all parameters have been resolved, along wit the information on whether the command finished successfully. On the next level, we can find QuNex summary (run) logs that show exact outgoing calls to external tools and pipelines for each session, along with the final command status. These summary logs also include errors that might have prevented the command from executing, such as missing parameters or mandatory input data. The most detailed logs are the command logs, which represent the exact console output of executed commands. For example, executing `hcp_freesurfer` produces the log identical to that produced by the HCP FreeSurfer pipeline. You can find examples of the generated logs for all three levels of granularity in the Supplementary Material document accompanying this manuscript.

The examples above demonstrate the general logic and usage of QuNex recipes through simplified placeholders. Practical examples applying recipes to actual MRI datasets are presented in the Results section.

### 2.3 Internal execution and workflow management

When the `run_recipe` command is invoked, QuNex parses the specified YAML file and constructs an internal representation of the workflow. The contents of the YAML file are automatically validated during parsing, and the process will abort if there are issues with the file’s formatting. This means that no processing will happen, and the user can correct the mistake and rerun the recipe without any downtime. Next, each command block is interpreted in sequence, with parameters resolved according to the hierarchical precedence rules described above. This parsing step ensures that all values defined in `global_parameters`, recipe-level parameters, and command-level parameters are consolidated into a unified parameter space. If dynamic substitution variables (defined via the mustache notation) are detected, they are resolved at runtime by querying the current environment.

After parsing, the recipe is executed through the QuNex workflow engine. Depending on the specified configuration and available computing resources, processing steps are launched sequentially or concurrently. Each command is executed within the containerised QuNex environment to ensure consistent access to dependencies and software versions. For computational clusters, the `run_recipe` framework supports native integration with SLURM, PBS and GridEngine scheduling systems. When executed on a high-performance computing (HPC) system, QuNex automatically generates scheduler submission scripts and manages dependencies between jobs, enabling the parallel or batched execution of independent steps.

Each executed command produces structured logs that capture the exact command line invocation, resolved parameter values, execution time, and return status. These logs are stored in a human-readable format to facilitate provenance tracking and post hoc quality control. Upon completion, a summary report is generated, indicating whether all steps completed successfully or if any command terminated with an error. The main validation mechanism for establishing whether a command has finished successfully is exit code of the command (the industry standard is that 0 denotes successful completion, and any other number denotes failure). If a command fails, the recipe will terminate, and execution can be resumed from the last successful step without re-running completed portions of the workflow. This can be done in two ways. The first way uses the `-steps` parameter, which represents a comma-separated list of commands to execute. The second way uses the `-startwith` parameter, where the user lists the name of the command to be used as the starting point within a recipe. When this option is used, preceding commands in the recipe are ignored. This checkpointing mechanism ensures efficiency and robustness in large-scale processing tasks. Note that the QuNex command validation is more sophisticated as it checks not only the exit code, but also the presence of mandatory outputs.

In addition to logging and error handling, the recipe framework records the QuNex container version, execution date, and system environment variables used during runtime. These records provide full provenance metadata, allowing complete replication of the processing environment and facilitating formal reproducibility audits.

Overall, the internal workflow management system enables QuNex recipes to function as self-contained, auditable processing specifications. By combining declarative workflow definitions with containerized execution and automated provenance tracking, QuNex ensures transparency, reproducibility, and scalability across a wide range of computational infrastructures.

### 2.4 XNAT integration

The Extensible Neuroimaging Archive Toolkit (XNAT; Marcus et al. 34) is an open-source imaging informatics platform developed by the Neuroinformatics Research Group at Washington University in St. Louis. XNAT provides a flexible and extensible environment for organizing, storing, and processing neuroimaging data, and supports execution of containerized workflows through its Docker and Singularity/Apptainer integration frameworks. The new recipe-based functionality in QuNex extends this compatibility, enabling streamlined and reproducible data processing directly within XNAT.

There are several advantages to executing QuNex via XNAT. In QuNex, MR sessions are organized in a flat directory structure, allowing efficient concurrent processing with minimal internal parsing. In contrast, XNAT organizes data hierarchically, with sessions nested under subjects and projects. While this structure facilitates data management, it requires that processing be launched on a session-by-session basis. For studies with many sessions, this approach can become inefficient, as the number of individual command calls grows linearly with dataset size. The recipe framework mitigates this limitation by bundling multiple processing and analysis commands into a single executable specification, thereby reducing the number of user interactions required to initiate large-scale processing and minimizing administrative overhead.

Another advantage of running QuNex through XNAT lies in its access control and user management capabilities. Within XNAT, administrators can restrict which commands are available and what data can be accessed, ensuring compliance with project-specific data governance policies. Traditionally, this required administrators to predefine and maintain multiple command configurations – an approach that is both time-consuming and privileges-dependent. By adopting recipe-based processing, administrators now need only authorize a single `run_recipe` command. Project supervisors and advanced users can then define project-specific recipes containing customized commands, parameters, and checkpoints, all without requiring elevated administrative permissions. This design significantly accelerates user onboarding and reduces the risk of executing incorrect or unauthorized commands.

Furtermore, XNAT employs a working-directory model, in which archived data are temporarily copied to a processing workspace before execution. Upon successful completion, results are copied back into the archive; failed or incomplete processing runs can be safely discarded, preventing contamination of validated datasets. Although this model enhances data integrity, it can become inefficient for large studies, as each command invocation requires full data transfer. This also complicates iterative reprocessing, since intermediate results may be repeatedly reloaded. The recipe framework addresses this issue by automatically creating *checkpoints* – snapshots of the working directory before and after each processing step. These checkpoints can be recalled during subsequent runs, allowing selective re-execution of failed or updated steps without redundant data transfer. This approach greatly improves both computational efficiency and reproducibility in large-scale or multi-stage analyses.

QuNex provides a wide range of configurable commands and parameters, which, while powerful, may be challenging for new users. Traditionally, neuroimaging processing and analysis is a computationally complex task executed on highperformance computing systems using containers (e.g., Singularity, Docker) and schedulers (e.g., SLURM, PBS, GridEngine, etc.). Many neuroscience researchers lack a strong technical background and are overwhelmed by the complexities that these technologies present. QuNex simplifies many of these steps through its own container and a specialised `qunex_container` script for interacting with it, as well as through a simplified, semi-automated scheduling system (see (1), for details). However, users still need to connect to a remote system and interact with QuNex via a command-line interface. Using XNAT significantly alleviates the problems arising through these additional layers of complexity.

Overall, integrating QuNex recipes with XNAT provides a unified and efficient mechanism for controlled, reproducible, and scalable processing of neuroimaging data, leveraging the strengths of both platforms in data organization, user-friendliness and workflow transparency.

## 3. Results

The Results section illustrates the practical use and reproducibility of the QuNex recipe framework. We first describe the official repository accompanying this manuscript, which contains all materials required to reproduce the examples presented below. In addition, the repository provides an extensive collection of validated recipes that have already been successfully applied in several large-scale and clinical neuroimaging research projects.

We then showcase two representative use cases that demonstrate the flexibility of the recipe framework in real research settings. The first example (Listing 2) showcases an end-to-end workflow beginning with raw MRI data in the DICOM format. The data are converted to NIfTI, onboarded into a QuNex study, processed using the HCP Minimal Preprocessing Pipelines (8), and subsequently analysed for functional connectivity. This use case illustrates how a complete data-processing and analysis pipeline can be encoded and executed through a single recipe, ensuring transparency and full reproducibility.

The second example (Listing 3) highlights the integration of external data sources and specialized diffusion analysis tools within the same framework. Here, preprocessed HCP data (35) are downloaded via an external script, onboarded into a QuNex study, and analysed using FSL’s DTIFit (36) followed by Neurite Orientation Dispersion and Density Imaging (NODDI) microstructure modelling (37, 38). Together, these examples demonstrate how QuNex recipes can support both end-to-end and modular analytic workflows while maintaining complete provenance and reproducibility. Unless otherwise specified, all examples assume that the official QuNex recipe repository is located at `/data/qunex_run_recipe` and that QuNex studies are stored under `/data/studies`.

### 3.1 The official QuNex recipe library

The official repository associated with this manuscript (https://github.com/ULJ-Yale/qunex_run_recipe) is publicly accessible without registration and provides a version-controlled library of QuNex recipes, together with all the materials needed to reproduce the examples presented herein. The repository is organised into the following subfolders:

- `example_1` – materials, scripts, and configuration files for the first example described in this manuscript;
- `example_2` – materials, scripts, and configuration files for the second example;
- `recipes` – the main QuNex recipe library containing validated workflows for a variety of processing and analysis scenarios.

The recipe library currently includes a growing collection of tested workflows that have been applied across multiple research projects. These recipes cover a diverse range of use cases, including processing of HCP and HCP-style datasets, legacy data lacking T2-weighted images, imports from DICOM and BIDS formats, longitudinal analyses, diffusion modelling, and functional connectivity workflows.

The repository has been designed as a community resource that will evolve alongside the QuNex platform. Researchers are invited to contribute new or modified recipes to the library by submitting pull requests to the repository. All submissions are reviewed to ensure compatibility and reproducibility before being integrated into the public collection. External scripts can be added to the pull request and will then be added to the newly created folder for the uploaded recipe. This open model supports the transparent sharing of the best analysis practices and accelerates the dissemination of standardised, reproducible neuroimaging workflows. Furthermore, we plan to update the software stack and QuNex functionalities to support use cases implemented in commonly encountered community external scripts.

### 3.2 A recipe file for end-to-end processing

The first example demonstrates an end-to-end workflow that converts raw MRI data in the DICOM format into preprocessed and analysis-ready outputs suitable for functional connectivity analyses. The recipe includes data onboarding, preprocessing using the HCP Minimal Preprocessing Pipelines (8), HCP ICAFix denoising (39) and HCP MSMAll surface matching (40, 41), running additional functional connectivity preprocessing steps, and computing Global Brain Connectivity (GBC) maps on a combined surface and volume representation across a single resting-state BOLD image. GBC provides a data-driven functional MRI measure that quantifies the average strength of connectivity between each voxel or cortical vertex and all other regions of the brain, providing an index of global network integration (42). This example illustrates how a complete, reproducible processing and analysis workflow can be encapsulated in a single recipe file, enabling transparent documentation and one-command execution.

**Listing 2.**
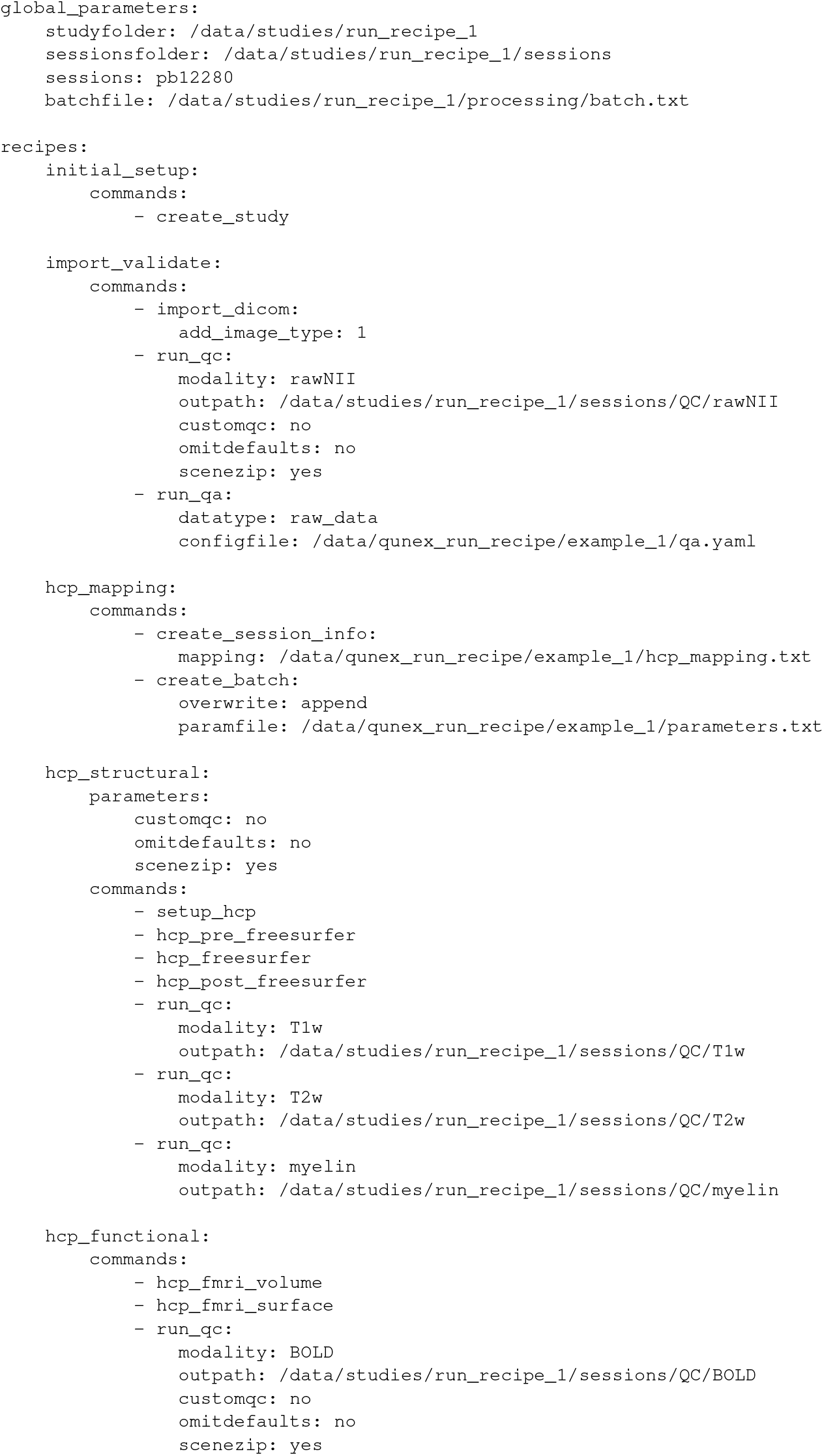
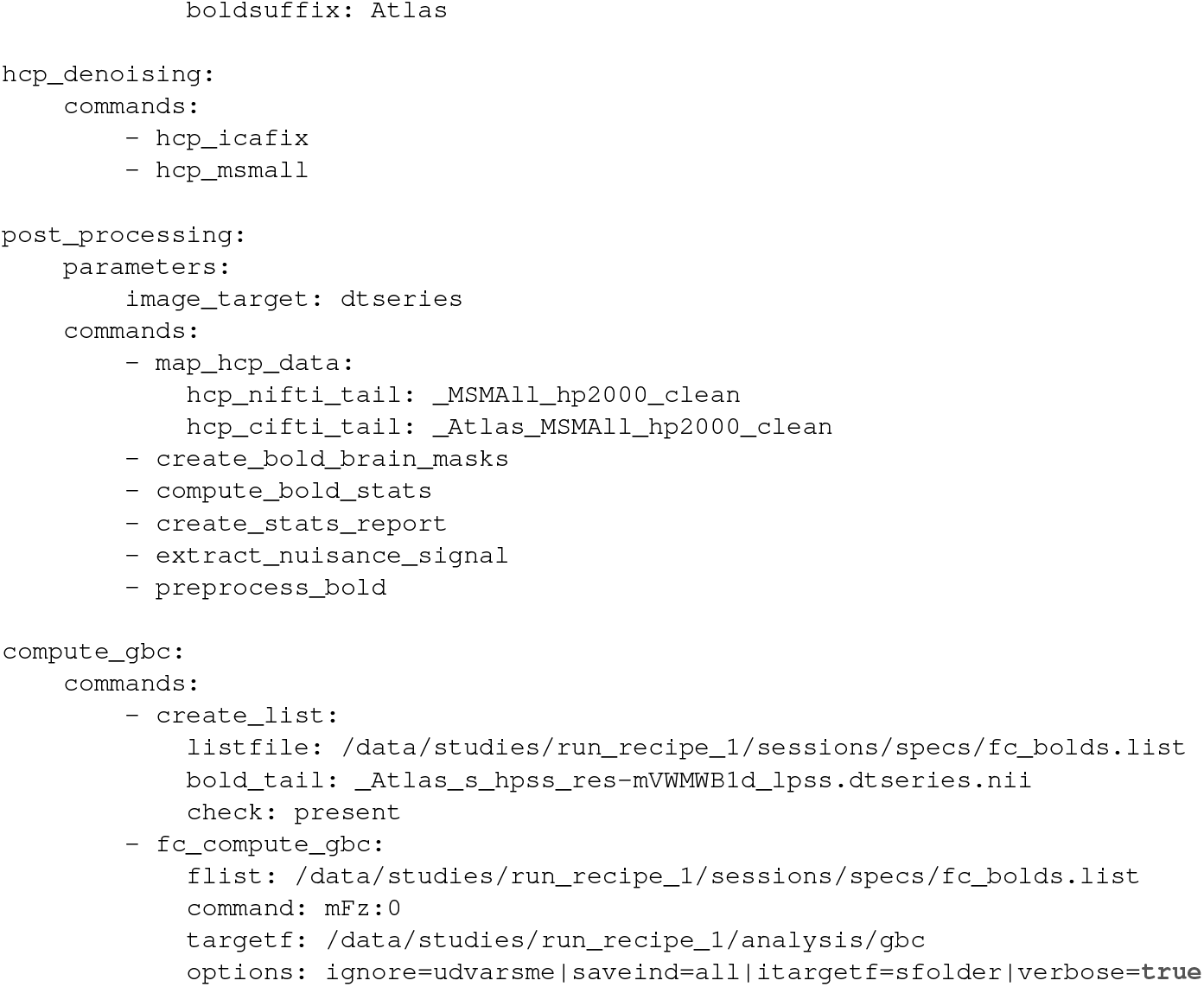
A recipe for end-to-end processing.

For illustration purposes, the example specifies a single session (`pb12280`); however, in typical studies, multiple sessions would be processed, usually by invoking the recipe through a command-line call rather than by manually editing the recipe file. The example recipe is composed of seven distinct sub-recipes, listed in the order in which they should be executed. This structure creates purposeful breakpoints at which users can verify that each stage has completed successfully and that intermediate outputs meet expected quality standards. It also establishes a modular system that facilitates the addition of new recipes for extended processing methods – for example performing seed-based functional connectivity analyses.

Here, we demonstrate how QuNex can be used in conjunction with QuNex recipes to execute all preparatory steps, processing and analyses in a reproducible manner, and present the full workflow and all parameters in a transparent, human-readable way. For comparison, the Supplementary Material document for this manuscript contains the same example, but without the QuNex recipes framework, using the traditional command invocation mechanism instead. That approach is significantly longer, more tedious, harder to read and comprehend as well as prone to errors.

The first recipe, `initial_setup`, is executed once per study or dataset and initializes the folder hierarchy required by QuNex. The remaining recipes are executed for each MRI session and can be grouped into preprocessing and postprocessing stages.

The preprocessing stage includes the `import_validate`, `hcp_mapping`, `hcp_structural`, and `hcp_functional` recipes. These steps handle the import of raw DICOM files, validation of acquisition parameters and scan mapping, and execution of the HCP Minimal Preprocessing Pipelines. During preprocessing, QuNex automatically generates quality control (QC) snapshots and Connectome Workbench (43) `.scene` files, enabling both rapid and detailed visual inspections of intermediate results. These QC outputs should be reviewed at each breakpoint before continuing with subsequent stages. Apart from the parameters specified in the QuNex batch file (see (1) and https://qunex.readthedocs.io), these preprocessing steps are largely standardized across studies. In most cases, only minor adjustments – such as modifying the import command for datasets organized in the BIDS format – are required.

The final two recipes, `post_processing` and `calculate_gbc`, perform mapping preprocessed data to the QuNex folder structure, denoising and analyses. When mapping the data from HCP to QuNex folder structure for analyses, users can specify which data to use by using the `hcp_nifti_tail` and `hcp_cifti_tail` parameters. In our case, we will use minimally preprocessed data, that has been also denoised and sufrace matched on top. Denoising includes regression of nuisance signals derived from ventricular, white matter, and whole-brain masks, together with motion parameters, from the BOLD time series. The resulting residual time series are then used to compute GBC maps. These steps often require project-specific adaptation, which highlights the benefit of the modular recipe design.

This modular approach allows users to customize and extend the workflow while maintaining full transparency and provenance of all analytic steps.

### 3.3 A recipe for starting with existing data

The second example illustrates how QuNex recipes can be used to build upon existing datasets by executing additional diffusion processing steps on already preprocessed data. This use case originates from one of our research projects, in which HCP data were downloaded from a public S3 archive, imported into QuNex, and processed using FSL’s DTIFit (36) and Neurite Orientation Dispersion and Density Imaging (NODDI) microstructure modeling (37, 38).

**Listing 3.**
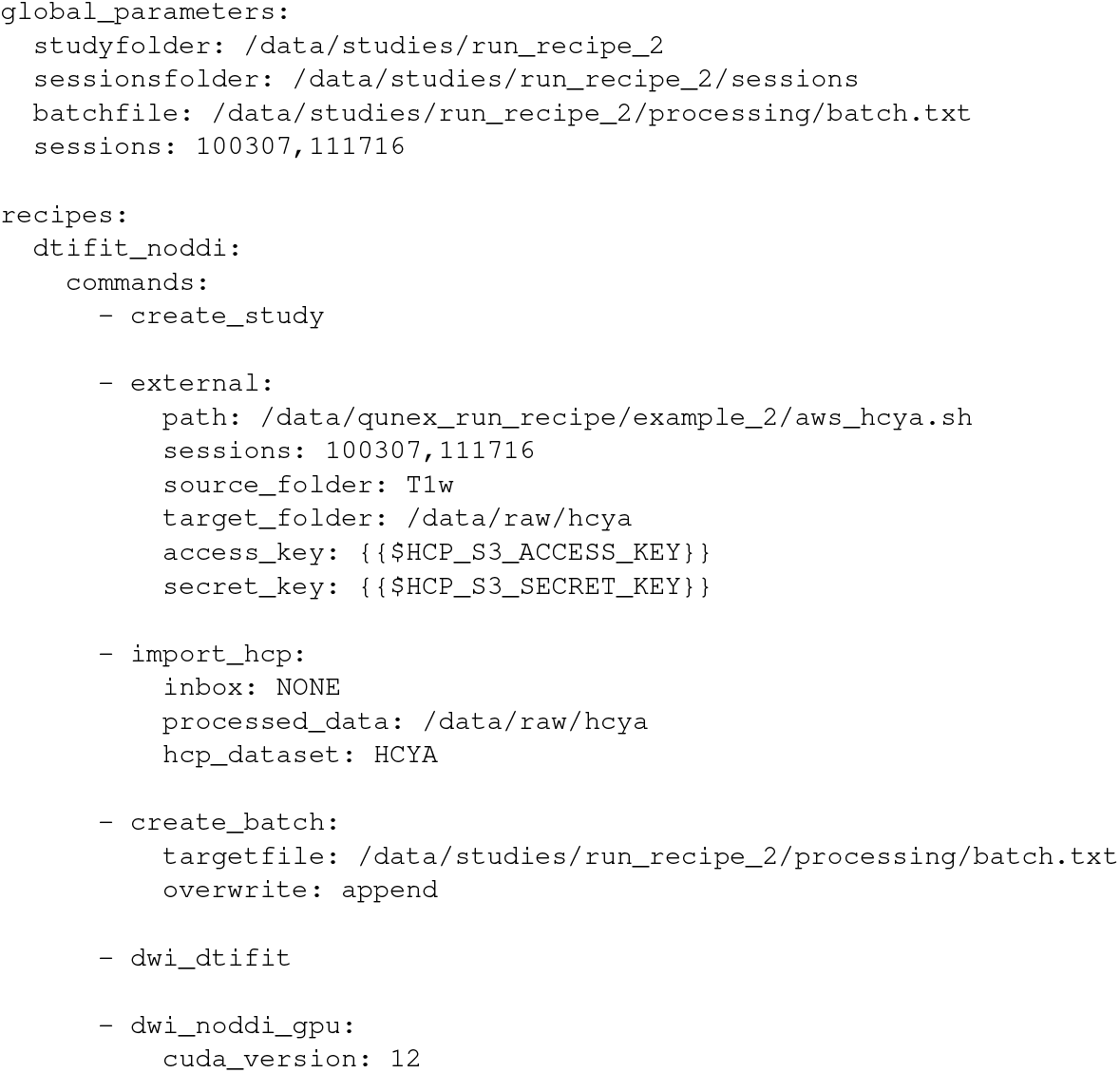
A recipe for DWI NODDI processing of preprocessed HCP data.

This recipe executes six sequential steps across two sessions (HCP Young Adults participants `100307` and `111716`). Two sessions are shown here for brevity; however, the same workflow can be readily scaled to thousand sessions or more. By default, QuNex processes sessions sequentially. Parallel processing can be achieved either by launching the recipe simultaneously for multiple sessions or by using a scheduler such as SLURM (https://slurm.schedmd.com/overview.html). When a scheduler is employed, QuNex automatically generates one job per session, enabling efficient distributed execution. For details about the QuNex scheduling framework, please refer to the official QuNex manuscript (1) and documentation https://qunex.readthedocs.io. The example below demonstrates how the recipe can be executed with GPU support through a scheduler:

~~~
qunex_container run_recipe \
  --recipe_file=“/data/qunex_run_recipe/example_2/dtifit_noddi.yaml” \
  --recipe=“dtifit_noddi” \
  --nv \
  --scheduler=“SLURM,time=01-00,mem=32G,gres=gpus:1,jobname=qx” \
  --container=“/data/qx_containers/qunex_suite-1.4.3.sif”
~~~

The recipe begins with `create_study`, which initializes a new QuNex study directory and folder structure. The following command, an external script (`aws_hcya.sh`), downloads the required data from the HCP Amazon S3 bucket (for access instructions, see the HCP Wiki at https://wiki.humanconnectome.org). The next step, `import_hcp`, onboards the downloaded data into the QuNex study, followed by `create_batch`, which generates a batch file containing session-specific metadata and processing parameters. For a detailed description of batch files and their purpose, please refer to the official QuNex manuscript (1). These initial steps can be categorized as data onboarding and preparation.

The final two commands, `dwi_dtifit` and `dwi_noddi_gpu`, perform the required diffusion processing. The first fits diffusion tensors using FSL’s DTIFit, while the second estimates NODDI parameters using GPU acceleration (indicated by the `_gpu` suffix). To enable GPU execution, the `-nv` flag must be specified in the `qunex_container` call in order to properly load the GPU libraries inside the container, and a GPU must be requested through the scheduler configuration parameter (e.g., `gres=gpus:1`).

Although only two sessions are shown here, this approach can be scaled to large datasets with minimal modification. In our application, this recipe was used to process the entire HCP Aging dataset – over 1,000 sessions – by executing a single recipe command that automatically handled downloading, onboarding, and diffusion processing. This example demonstrates the scalability of the QuNex recipe framework and its capacity to integrate public datasets, external scripts, and high-performance computing resources into a fully reproducible workflow.

### 3.4 Summary

Together, the two examples demonstrate the flexibility, scalability, and transparency of the QuNex recipe framework. The first example shows that complete end-to-end processing – from raw DICOM data to fully preprocessed and analysed outputs – can be achieved reproducibly through a single, human-readable recipe. The second example illustrates how the same framework can integrate preprocessed data, external scripts, and high-performance computing resources to execute specialized analyses at scale. Across both use cases, all parameters, commands, and software versions are explicitly recorded within the recipe and container environment, ensuring full provenance and facilitating replication of results on any compatible system.

By combining containerized execution, structured parametrization, and a declarative workflow specification, QuNex recipes unify what have traditionally been fragmented stages of neuroimaging analysis. This approach enables researchers to move seamlessly between local, institutional, and cloud computing environments, while maintaining a fully documented and reproducible record of their analytic workflow. In doing so, the recipe framework extends the QuNex platform beyond preprocessing toward a general-purpose system for transparent, scalable, and shareable neuroimaging pipelines.

### 3.5 Limitations

Although QuNex aims to simplify neuroimaging workflows at every stage, certain complexities are unavoidable due to the nature of the problem (e.g. remote use of specialised servers, command line interface, containers and schedulers). While XNAT can alleviate some of these issues, it is not widely used by research labs and requires the installation and deployment of additional software. Furthermore, the QuNex container is large (20 GB) and requires reasonably powerful hardware for local deployment, with ongoing expansion further increasing hardware demands.

One of the key limitations at present is suboptimal scheduling. Currently, the entire QuNex recipe is scheduled as a single job. However, if a study includes multiple sessions, we can use QuNex’s scheduling framework to create one job per session. This would result in massively parallel execution across a large number of sessions within a study. One of our future development tasks is to optimize this by decomposing recipes into sub-tasks, with each sub-task representing a scheduled job. To achieve this, we need to create an execution graph (a directed acyclic graph, DAG) that outlines the dependencies between these tasks and can serve as the basis for optimal scheduling. For example, if we wanted to run HCP structural and functional processing on a single session with T1w, T2w and four rest BOLDs, this would be represented in the execution graphs. In the current QuNex recipe framework this would run through a single scheduler job, with parallelism in functional processing achieved via multiprocessing within that job. However, this approach is suboptimal when we there are many BOLDs since it is not possible to process them all in parallel within a single job/node due to high computational demands of the task. The DAG approach would decompose the recipe into smaller tasks (e.g. processing of a single BOLD) and would spawn jobs for these tasks across many nodes, resulting in optimal processing. In this case, users will also be able to specify scheduling resources within a QuNex recipe at a command level.

## 4. Conclusion

QuNex recipes represent a significant advancement toward reproducible, transparent, and scalable processing of neuroimaging data. By providing a standardized, human-readable, and machine-executable format for defining analysis workflows, the framework bridges the long-standing gap between flexibility and reproducibility – two often competing objectives in neuroimaging research.

Through practical examples, we demonstrated how the recipe framework enables complete end-to-end analyses, from onboarding raw DICOM data to advanced connectivity and diffusion modeling, all within a single YAML-defined workflow. The framework seamlessly integrates containerized QuNex commands with external scripts and analytical tools, ensuring full interoperability between existing neuroimaging software and custom methods. Once accompanied by its recipe file, QuNex version, and input data, any study can be fully reproduced through a single command-line execution, providing an unprecedented level of reproducibility and transparency.

In addition, QuNex recipes extend the functionality of the XNAT platform by simplifying administrative setup, improving workflow traceability, and reducing opportunities for human error. These features make large-scale neuroimaging pipelines more manageable, lowering the technical barrier for new users while preserving advanced configurability for expert users and system administrators.

Ultimately, the introduction of QuNex recipes has the potential to transform how researchers design, share, and replicate neuroimaging workflows. By encouraging the publication of fully reproducible, executable pipelines, the framework promotes cumulative scientific progress and strengthens confidence in neuroimaging findings. Moving forward, we envision a growing open repository of community-submitted recipes – each representing a transparent, executable record of scientific work – that will serve as a foundation for collaborative, standardized, and reproducible neuroimaging research.

## Data and Code Availability

The code and materials needed to reproduce the results and run the recipes can be found in the official open repository of this manuscript: https://github.com/ULJ-Yale/qunex_run_recipe.

QuNex is open source. To access the code, please register at https://qunex.yale.edu/registration. You can find a small dataset for testing QuNex functionality in the Quick Start section of the official documentation: https://qunex.readthedocs.io/en/latest/wiki/Overview/QuickStart.html.

## Author Contributions

JD: Conceptualization, Methodology, Software, Validation, Formal analysis, Investigation, Resources, Data Curation, Visualization, Funding acquisition, Writing – Original Draft, Writing – Review & Editing. AK: Software, Validation, Investigation, Writing – Review & Editing. AM: Software, Validation, Investigation, Visualization, Writing – Review & Editing. SB: Software, Validation, Investigation, Data Curation, Writing – Original Draft, Writing – Review & Editing. LP: Software, Validation. ZT: Software, Validation. CF: Validation, Data Curation, Writing – Original Draft, Writing – Review & Editing. MH: Validation, Writing – Review & Editing. JLJ: Validation, Writing – Review & Editing. AA: Conceptualization, Supervision, Funding acquisition, Writing – Review & Editing. CK: Data Curation, Writing – Review & Editing. MS: Data Curation, Writing – Review & Editing. DD: Data Curation, Writing – Review & Editing. MG: Supervision, Funding acquisition, Writing – Review & Editing. LDN: Data Curation, Supervision, Funding acquisition, Writing – Review & Editing. YC: Data Curation, Supervision, Funding acquisition, Writing – Original Draft, Writing – Review & Editing. GR: Conceptualization, Methodology, Supervision, Funding acquisition, Writing – Review & Editing.

## Ethics Statement

This study did not involve the collection of new human or animal data. All example analyses are based solely on publicly available, fully anonymized datasets. Therefore, no institutional ethics approval was required.

## Supplementary material

### QuNex resources

This manuscript represents a major upgrade to the QuNex software suite (https://qunex.yale.edu). For a detailed description of QuNex, please consult the official QuNex manuscript (https://doi.org/10.3389/fninf.2023.1104508) and for additional details the official QuNex documentation (https://qunex.readthedocs.io). If you have any usage question, feature requests or issues with using QuNex, feel free to post on the official forum (https://forum.qunex.yale.edu).

### A command-by-command execution example

Code below showcases how the first QuNex recipe example from the manuscript (end-to-end processing) would look like using the “traditional” command-by-command processing workflow. As you can see, in terms of text this is significantly longer, this approach also introduces a lof of duplicate parameter definitions which is an unnecessary source of possible errors.

**Listing 4.**
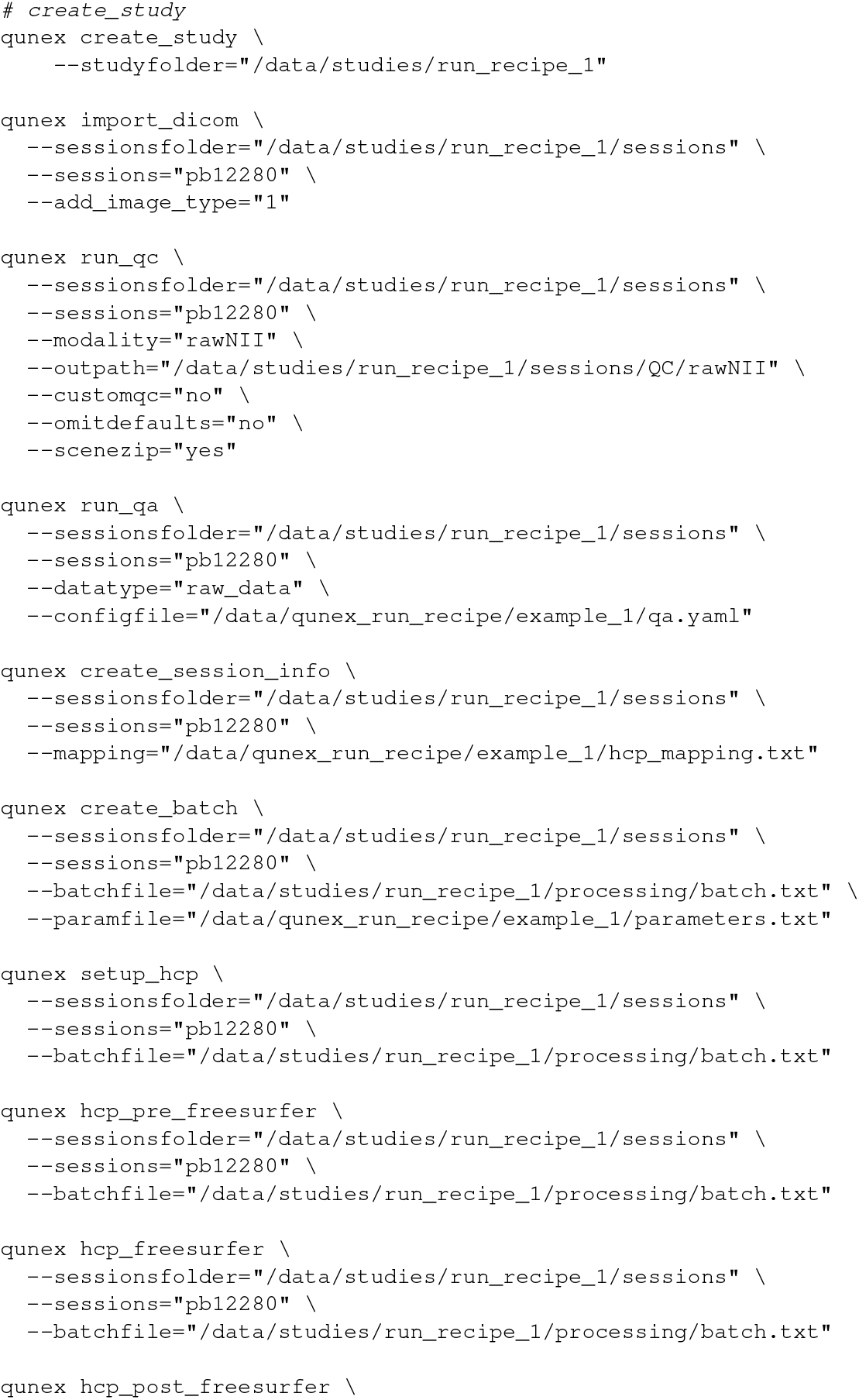
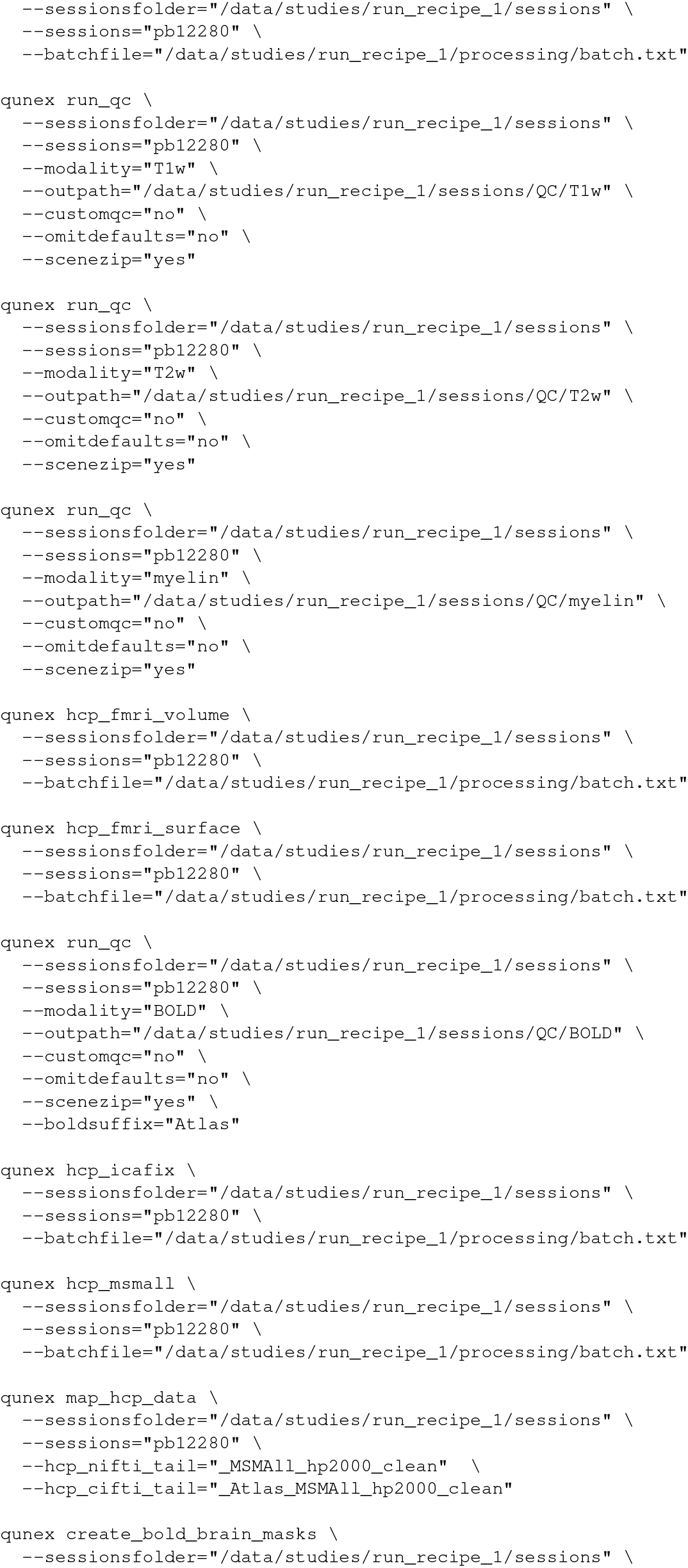
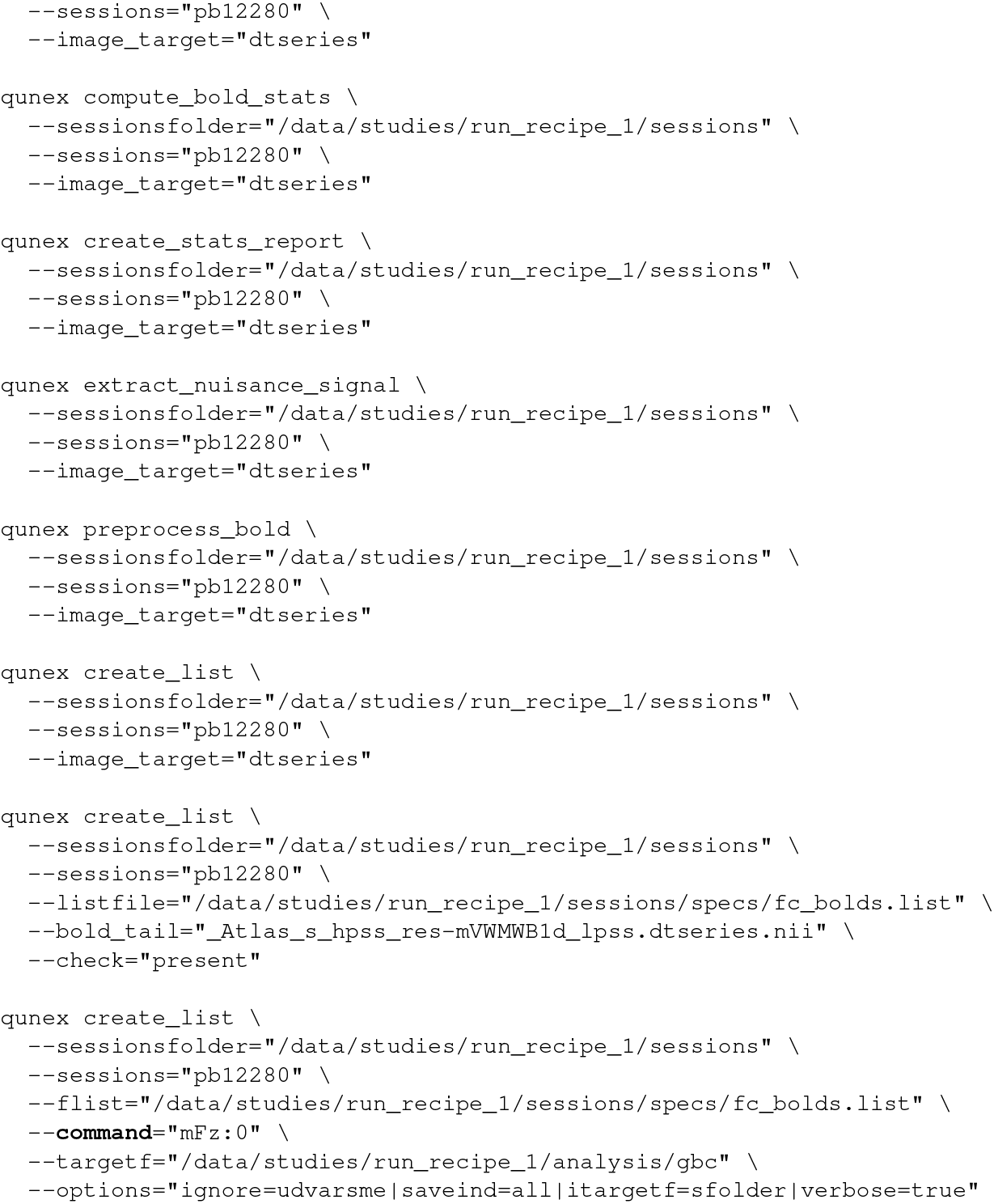
End-to-end processing example using QuNex commands.

### QuNex Recipes example logs

QuNex will generate logs at three different levels of granularity. Below are examples for each of these leves.

#### QuNex recipe logs

The top level logs are called recipe logs, these include a progress report on the highest level, the level of the whole recipe. Below is an example of such a log.

**Listing 5.**
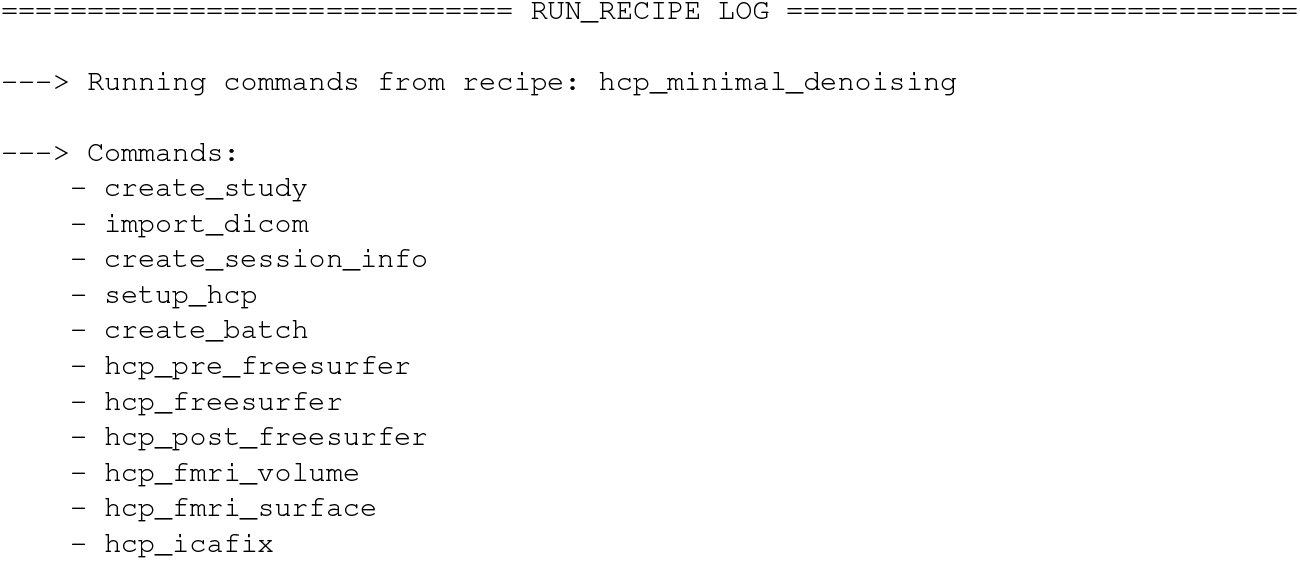
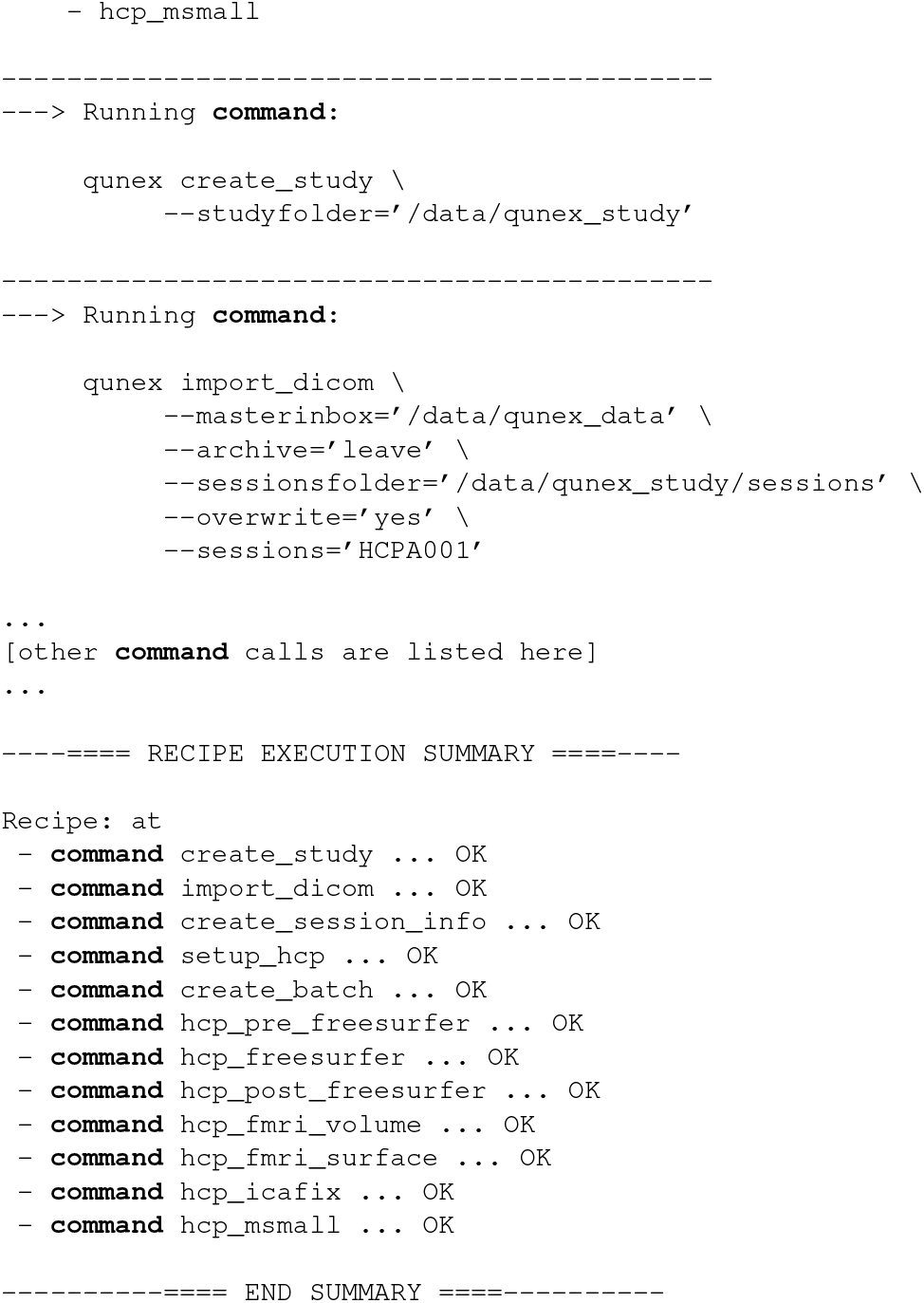
Example of the top-level recipe log.

#### QuNex summary (run) logs

One level below that are QuNex summary (run) logs, these do not include detailed printouts from commands but an overview of the command processing. Below you can find an example of this log for the `hcp_freesurfer` command.

**Listing 6.**
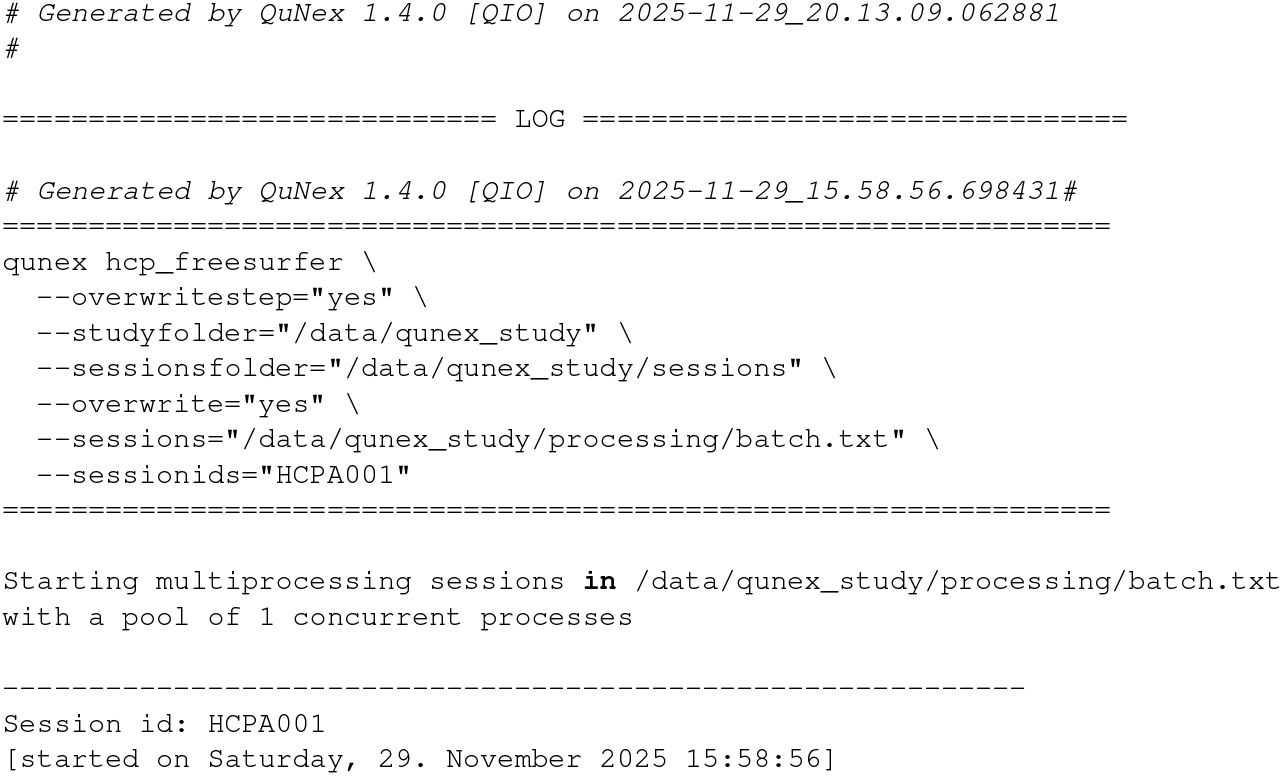
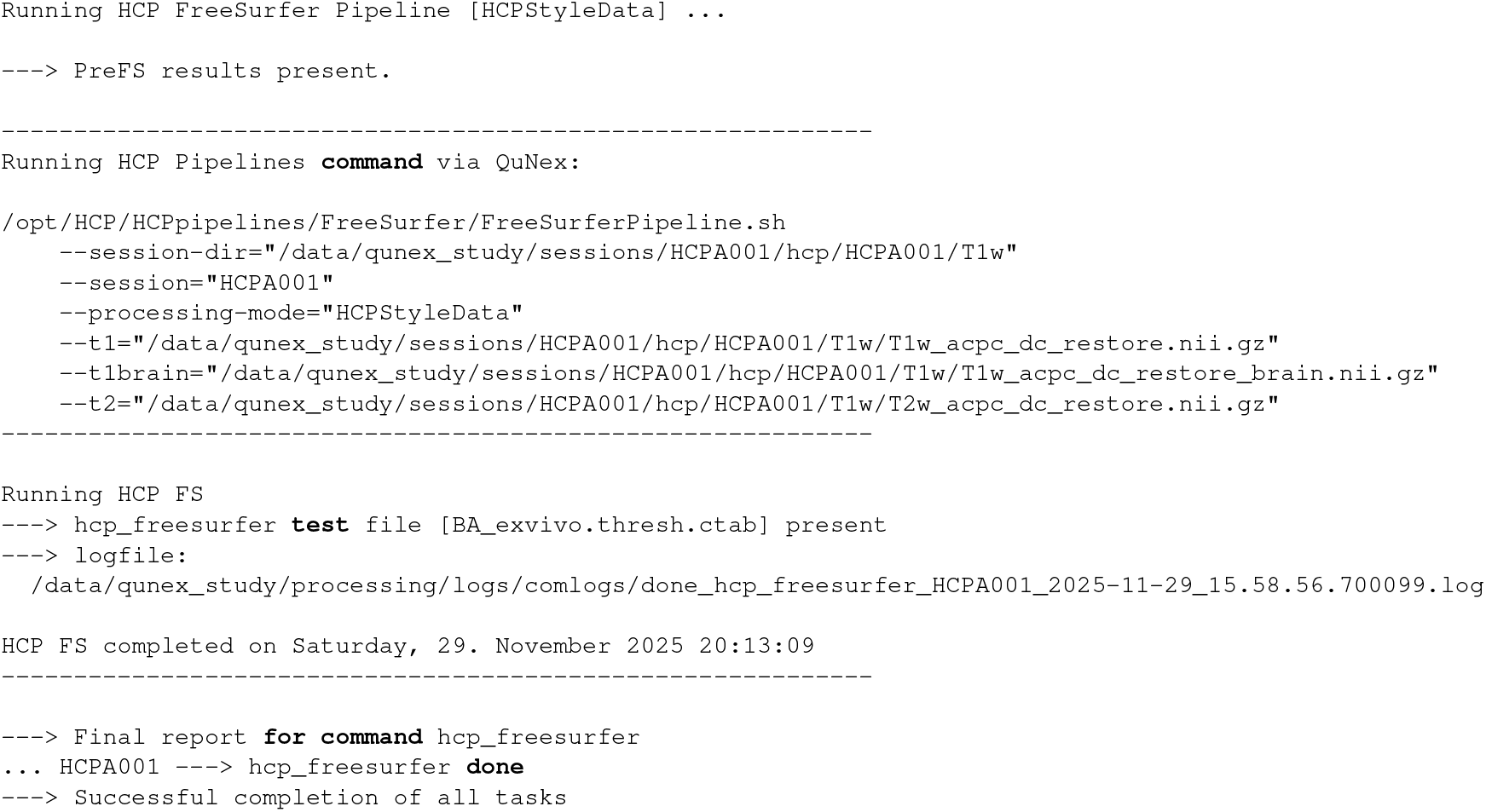
Example of the mid-level QuNex summary (run) log.

#### QuNex command (com) logs

At the lowest level are detailed command (com) logs. These include the complete printout of everything a commmand invocation prints into the OS console. Below is a shortened example for the above `hcp_freesurfer` command.

**Listing 7.**
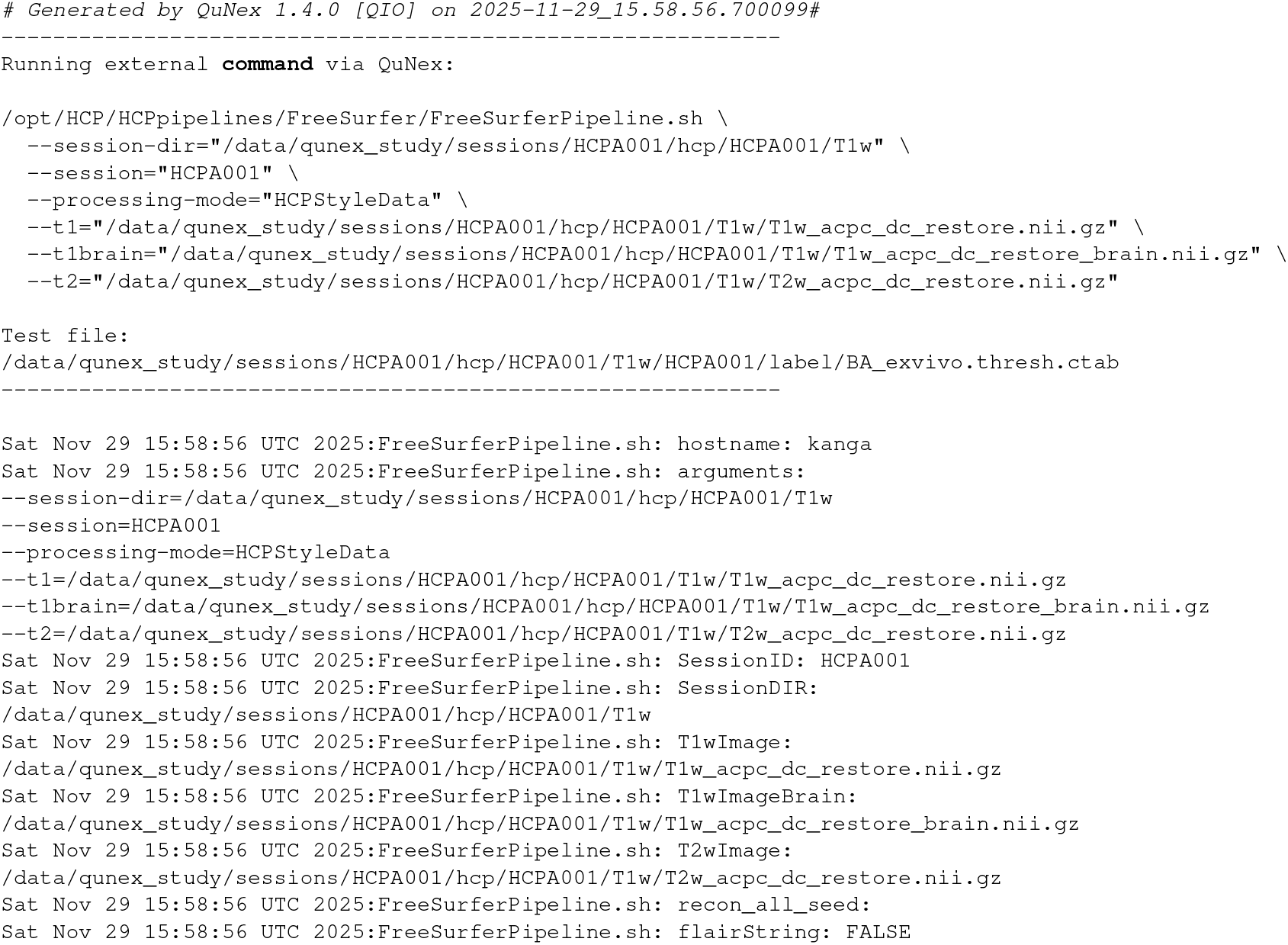
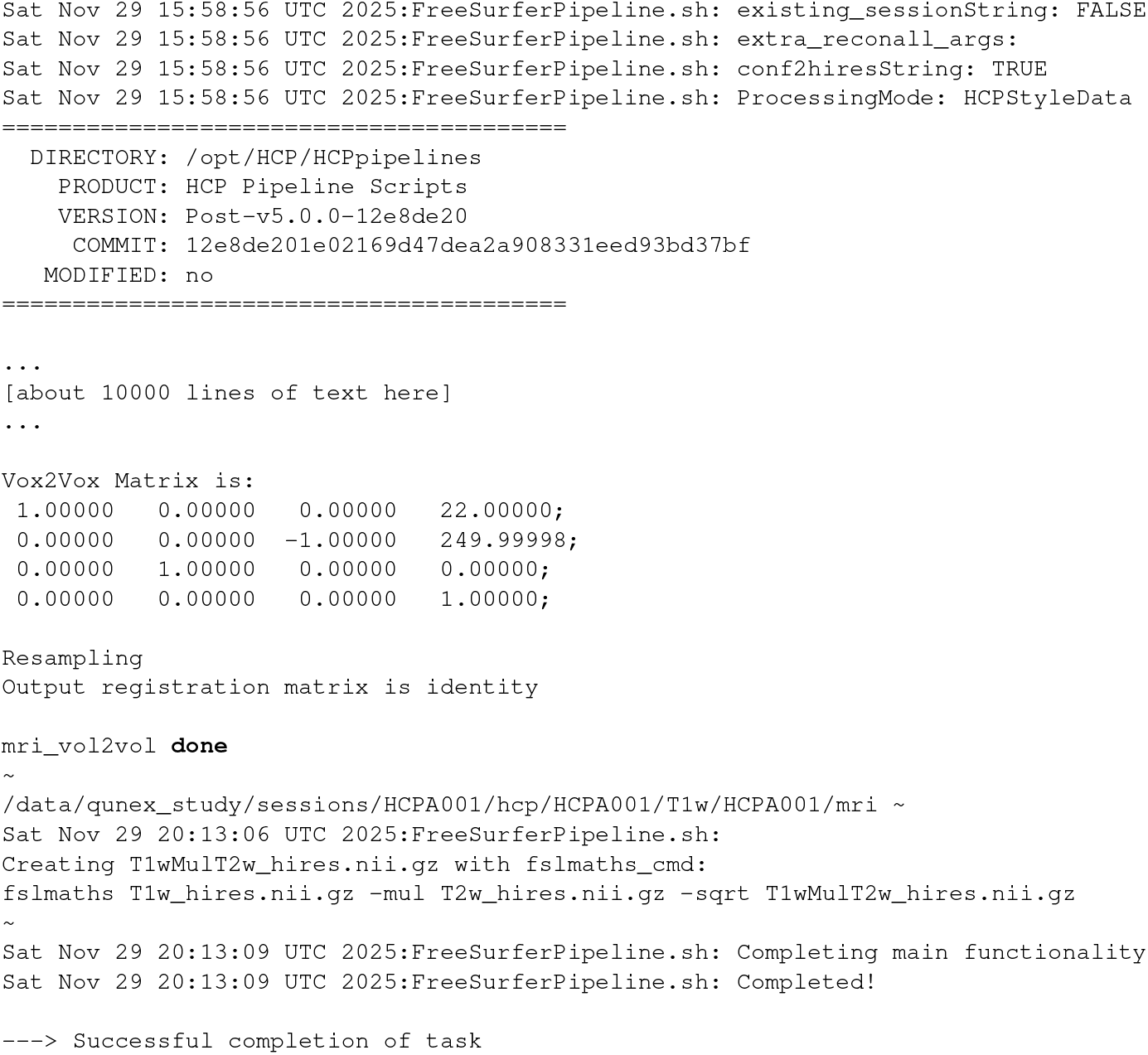
Example of the low-level detailed command (com) log.

